# *Chara canescens* part I: Oospore differentiation of parthenogenic and dioecious strains and salt-dependent oospore sizes

**DOI:** 10.1101/2024.03.26.586877

**Authors:** A. Holzhausen, R.E. Romanov, H. Schubert

## Abstract

*Chara canescens* Loisel. is one of two European species of the section Desvauxia R.D. Wood of the genus *Chara* L. Whereas most populations of *C. canescens* reproduce parthenogenetically, a few sites with sexual reproducing populations are known. Studies of European *C. canescens* oospore morphology led to open questions about the taxonomic status. Here we investigated nearly 1000 oospores from 16 European populations originated from plant release, sediments, germination experiments and herbaria and two Asian populations resulting in a regional determination key for studied populations as well as important database implications regarding origin, oospore plant position and equipment used. The impact of salinisation on oospore morphology were tested by artificial salt levels. The longest but smallest oospores were formed at 3 PSU, whereas the widest at 0.1 PSU and the shortest at 5 PSU. Basal width and shape, on the other hand, seem to be only affected by higher salt contents. This study contributes the lacking oospore-information for both reproduction modes of European of *C. canescens* – populations.

## 1. Introduction

Charophytes (Characeae) are one of the three lineages (Charophyceae, Zygnematophyceae and Coleochaetophyceae; Streptophyta) that form, together with land plants, the monophyletic group Phragmoplastophyta (Buschmann and Zachgo 2016). These macroscopic multicellular complex algae inhabit freshwater, as well as brackish and saline environments worldwide (Khan 1991). The relevance of Charophyceae for evolutionary and applied research areas, such as ecophysiology, paleoecology, ontogenesis, molecular and comparative genomics (Beilby 2019, Vicente et al. 2019, Holzhausen et al. 2022b, Quade et al. 2022) peaked in the establishment of the freshwater alga *Chara braunii* C.C. Gmel. 1826 as a model organism for water-land-transition of plants after publication of its draft genome (Nishiyama et al. 2018).

Within Characeae, the halotolerant species *Chara canescens* Loisel. (former *C. crinita* Wallroth 1815) is in three respects particular. First, it only can reproduce via oospores, secondly, it is the one of two European species of the section Desvauxia R.D. Wood and third, *C. canescens* is capable to reproduce via parthenogenesis, being a remarkable exception known among charophytes so far. This reproduction mode which is called apogamia in spermatophytes is a well-known phenomenon within flowering plants (Carman 1997, Hand and Koltunow 2014, Underwood et al. 2022) and described in detail for *C. canescens* by Ernst (1918) although the underlying molecular mechanisms that leads to geographic parthenogenesis of *C. canescens* are completely unknown so far (Schaible 2006).

Lessing seems to be the first who collected male plants from a bisexual population in the Caspian Sea and described them as a distinct species, *Chara karelinii* Lessing 1835 (Komarov Botanical Institute of the Russian Academy of Sciences, LE; Romanov 2021). Different opinions of leading botanist’s resulted in an incorporation of this dioecious population into the parthenogenetic *C. crinita* due to the absence of male species in field excursions over Central Europe (Braun 1857). Outside Europe, several monoecious and dioecious species from the section Desvauxia were described from Asia and North America. For example, T.F. Allen described monoecious *Chara evoluta* T. F. Allen and *C. hirsuta* T.F. Allen 1900 for North America, whereas monoecious *C. altaica* A. Braun in Braun and Nordstedt (1882), *C. sibirica* Migula 1904 were described from Siberia, monoecious *C. abnormiformis* Vilhelm 1928 from Central Asia, monoecious *C. piniformis* W.Q. Chen and F.S. Han in Chen et al. 1990 and *C.* F.S. Han 1964 from China, as well as dioecious Hollerb. in Hollerb. et Krassavina 1983 from Central Asia, *C.* Y.J. Ling et al. 1991, and *C. shanxiensis* Y.J. Ling 1985 from China. Male plants are known for *C. canescens* and *C. canescentiformis* but not found in case of *C. shanxiensis* and *C. pseudocanescens*. Still, until the discovery of *C. altaica*, found in two localities at the south of Eastern Europe (Romanov and Vishnyakov, in press), only one species of the section Desvauxia, i.e., *C. canescens*, was known from Europe. This species is represented in Europe mostly by parthenogenetic populations with a few sexually reproducing ones found in France, Italy (Sicily, Sardinia), Greece, Austria, and Serbia (Krause 1997, Nowak et al. 2019, Sabovljević et al. 2022). Outside Europe, sexually reproducing populations are reported from North Africa (Morocco), Central Asia, West Siberia, and China (Hollerbach and Krassavina 1983, Fushan and Yaoying 1994, Nowak et al. 2019, Romanov unpubl. data).

For Europe, the Austrian dioecious population is investigated in more detail. Decreased sodium chloride concentrations and increased concentrations of soda and sulphate are reported as decisive for their restriction (Krause 1997). Further, ecological conditions such as light irradiance and salinity do not have an impact on the occurrence of male *C. canescens* (Schaible and Schubert 2008).

The conservation of parthenogenic *C. canescens* strains require reliable taxonomic determination for both cases. If alga material is present, differentiation by use of *rbc*L and *atp*B genes can be used as successful solution (Schaible et al. 2009, Nowak et al. 2019). But parthenogenetically and sexually reproducing species are well known from temporary habitats in Spain, Austria, and Italy. The changes caused by climate change, such as longer and more frequent periods of drought, require the identification of oospores from sediments.

Taxonomical use of oospores is challenging (Holzhausen et al. 2023). One solution for a) identification of regional differences, b) equipment and storage-based differences and c) linkage between oospores and environmental data is the buildup of an worldwide oospore database (currently: Rostocker oospore-charophyte database, Anja Holzhausen) as recommended by Holzhausen et al. (2015) and described in Holzhausen et al. (*In Press*). This combines morphology data of oospores and alga, environmental data and few genetic analyses to assess those open fundamental questions and allow re-evaluation by storage of oospores and herbarium material.

The present study resolves the question of oospore differences by contribution of lacking oospore-information for both reproduction modes of European *C. canescens* – populations. The discovery of another member of the section Desvauxia in Europe raises the question of the origin of the parthenogenetic taxon.

## 2. Materials and methods

Nearly 1000 oospores from 16 European populations and one Asian population of *C. canescens* were analysed for quantitative and qualitative morphological parameters including the structure of the oospore wall. Table 1 lists all populations examined, 10 out of 16 were obtained from fresh plant material, oospores were harvested after release (Austria, Germany, France, Kazakhstan). Five populations were obtained from laboratory germination experiments (Austria, Germany, France, Spain), one from a sediment sample (Germany) and one from herbarium (France). Additionally, oospores of one *C. altaica* population from Kazakhstan was included (fresh material). For light microscopy (LM), oospores were placed on microscope slides with an adhesive surface and documented with a stereomicroscope in lateral, apical and basal view (Olympus SZX16 or Leica DM2500). Oospore colour, shape and appendixes were determined. Parameters such as oospore length, oospore width, fossa width, basal width and angle of striae to the longitudinal axis were measured using ImageJ (v. 1.50i).

**Table 1.**
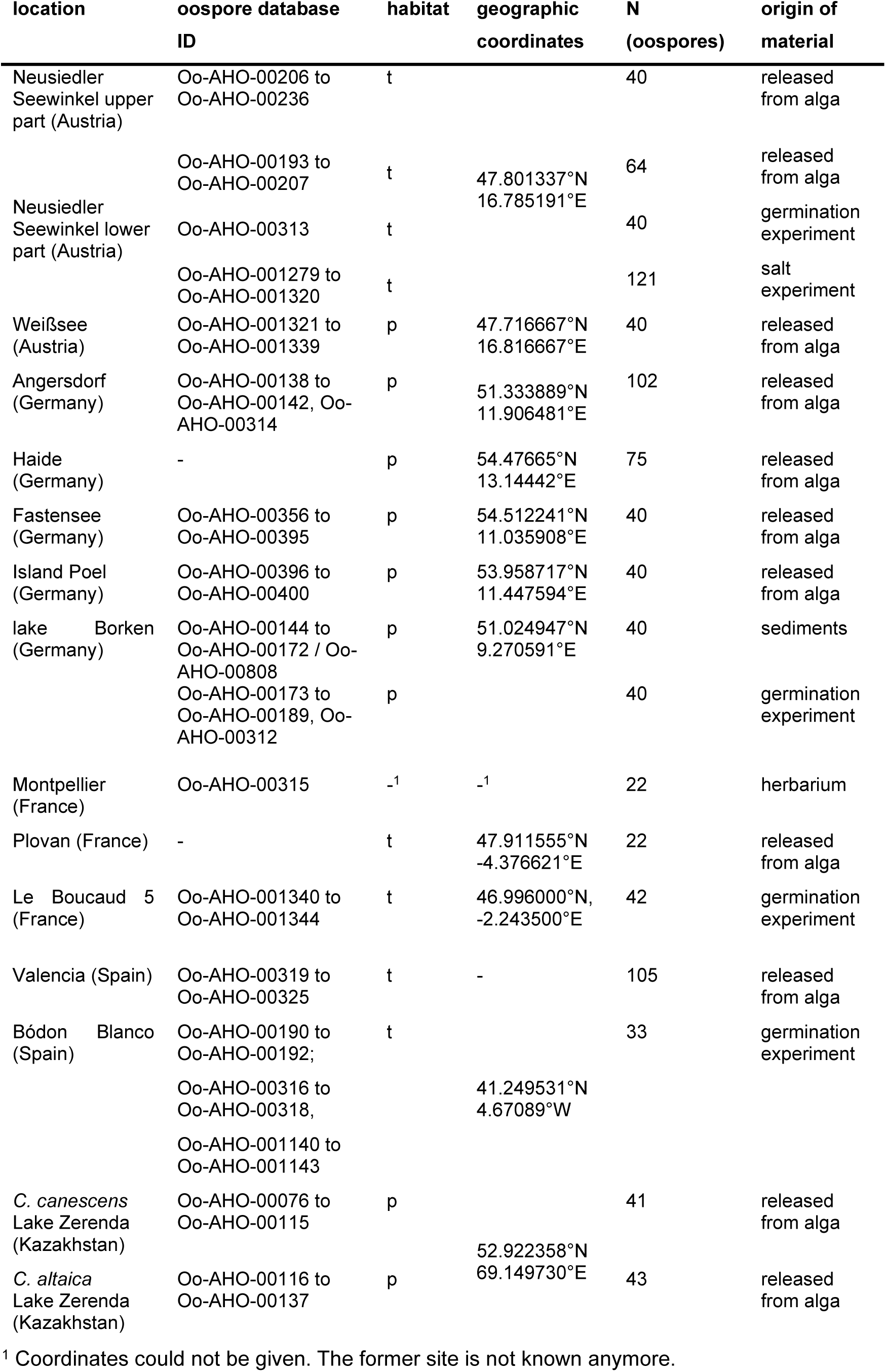
Summary of *C. canescens* and *C. altaica* populations used. Listed are the sampling site, habitat (temporary (t), permanent(p)), geographic coordinates (latitude, longitude), number of analysed oospores (N), the kind of material as well as the oospore-database ID (living material).

**Table 2.** Physico-chemical parameter of *C. canescens* sampling sites.

| Site | pH | salinity<br>(PSU) | conductivity<br>($\mu\text{S}/\text{cm}$) | oxygen<br>(mg/L) | T ($^{\circ}\text{C}$) |
| --- | --- | --- | --- | --- | --- |
| Angersdorf | - | 7 | 62.4 | - | 12.6 |
| lake Borken | 8.43 | 0.41 | 710 | 8.40 | - |
| Fastensee (Fehmarn) | 8.3 | 15.9 | 20120 | 7.81 | 7.9 |
| Haide | - | 9.7 | - | - | - |
| Poel | - | 13.5 | - | - | - |
| Neusiedler Seewinkel, lower part | 9.65 | 5.19 | 8710 | 8.96 | 15.8 |
| Neusiedler Seewinkel, upper part | 9.57 | 4.6 | 6960 | 8.46 | 15.7 |
| Weißsee | 8.63 | 1.32 | 2270 | 8.28 | 18.5 |

Scanning electron microscope analysis (SEM) of the oospore wall structure were done at the National History Museum London by M. T. Casanova. Oospores were dried by lyophilization and sputter-coated with gold but not treated with surface cleaning agents. SEM images of the surface of oospores and fossa walls were produced according to Wilbraham and Casanova (2019).

Statistical analyses were performed by using R (Team R 2022). Data were tested for normal distribution by Shapiro-Wilk test from the EnvStats package and for homogeneity of variance across groups with Levene’s test from car package (Fox and Weisberg 2019). ANOVA were performed for normal and homogeneous distributed data and Kruskal-Wallis rank sum tests for non-Gaussian (not normal) distributed data or data with heterogeneous variances. For data with significant ANOVA results Tukey’s HSD post-hoc tests were performed. In case of significant results from the Kruskal-Wallis tests multiple comparisons were done by Dunn’s test (Dinno 2017).

Box plot visualisation was conducted by the R packages ggplot2 and ggstatsplot (Wickham 2016, Patil 2021). Principal component analysis was performed and visualised using the R packages ggbiplot and corrplot (Wei et al. 2017).

## 3. Results

In a first step, morphological oospore differences possibly caused by their origin (oospores from sediments vs. oospores from habitat plants vs. in vitro germination experiments), the microscope used or the whorl number at which the oospores are grown were tested.

### Differences between oospores from sediments, the field and *in vitro* germination experiments

Oospore length (p.adj. = 5.33e-16), number of striae (p.adj. = 1.74e-2), colour (p.adj = 0.00387), mean basal width (p.adj. = 1.42e-8) and length/width ratio (p.adj. = 4.02e-7) of oospores from sediment samples of lake Borken differ significantly from oospores from in vitro germination experiments. These differences are mainly caused by broadened size ranges of oospores from sediment samples, except for the number of striae (Tab.3).

**Table 3.**
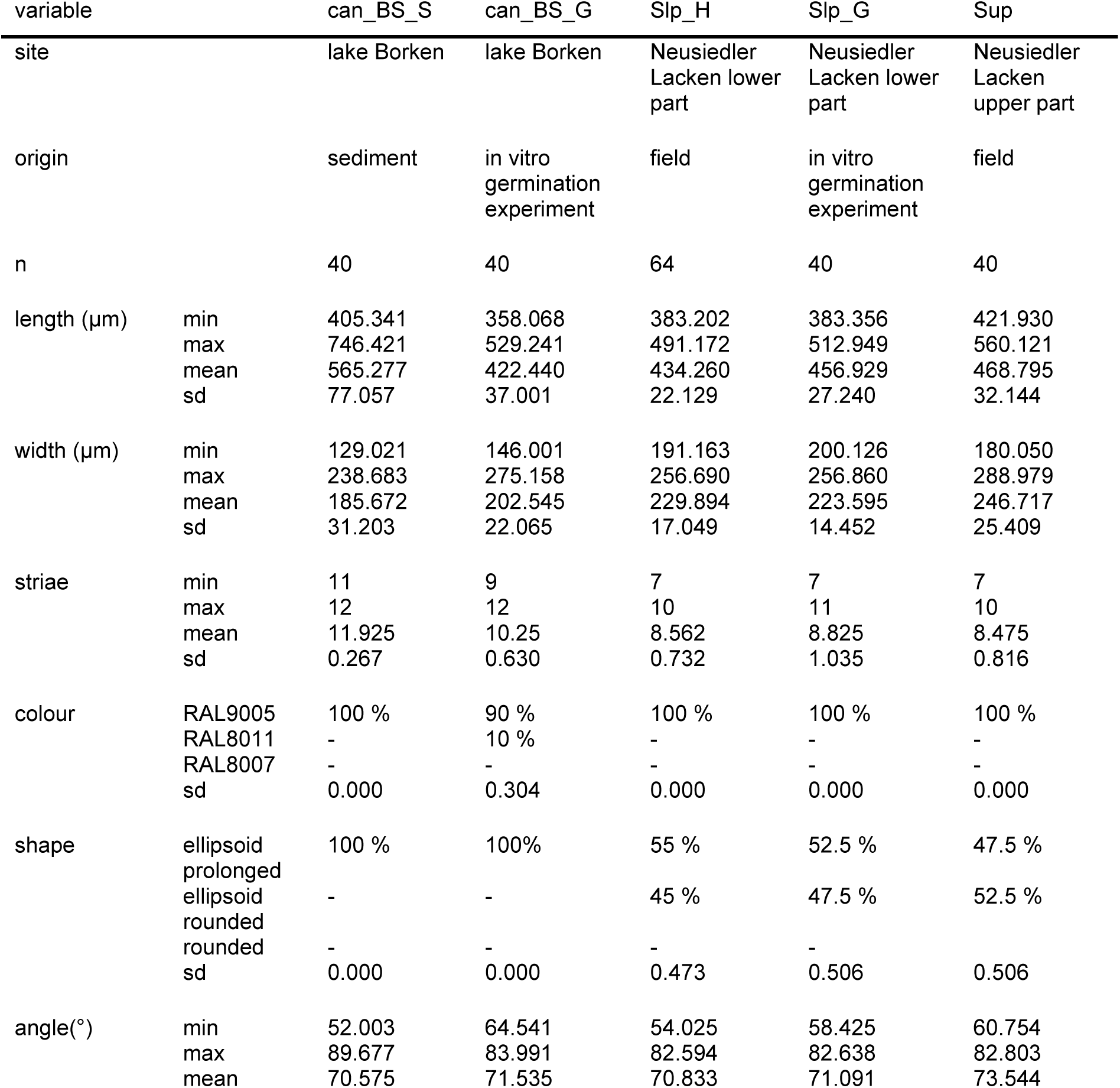
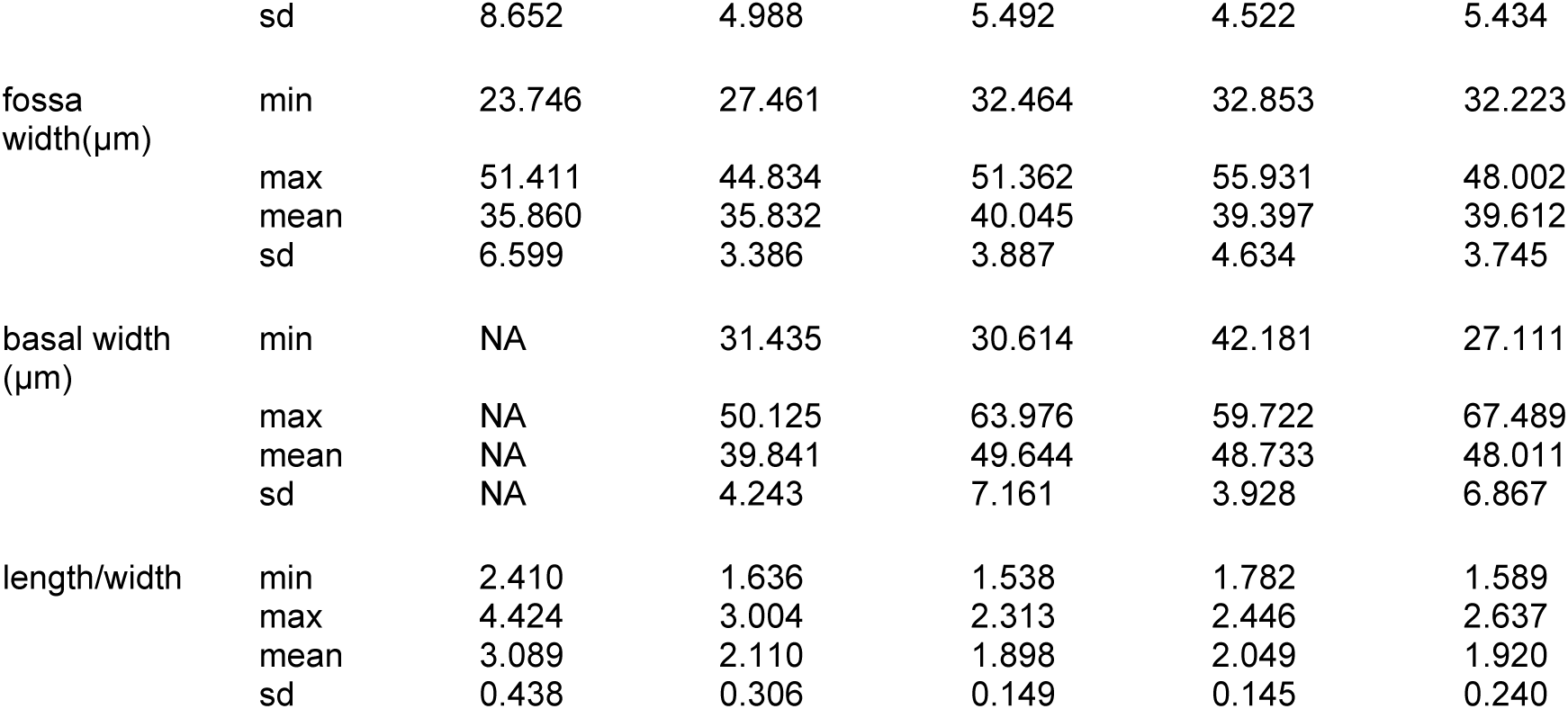
Overview of oospore characteristics of *C. canescens* populations from sediment samples, in vitro germination experiments and the field.

In contrast to this, no significant differences were found between oospores from field material and the corresponding in vitro germination experiment, except for oospore length (p.adj. = 2.732e-2) and the calculated length to width ratio (p.adj. = 8.75e-3). Despite small differences in size ranges, the calculated means of both oospore populations are consistent for all other oospore characters.

### Salt-dependent oospore differences

Oospores formed in medium with artificially increased salt content show clearly significant differences between the salt levels and to the field population (S-Tab.1).

For all characters, except the oospore colour (RAL9005), significant differences were found for at least two of the three salt levels (0.1 PSU, 3 PSU, 5 PSU, S-Tab.1) and the corresponding reference group (Can_Slp_G). Salinisation has the greatest effect on oospore length, oospore width, fossa width and number of striae. Figure 1 showed boxplots for these parameters and all groups. The longest but smallest oospores were formed at 3 PSU, whereas the widest at 0.1 PSU and the shortest at 5 PSU (Tab. 4). Basal width and shape, on the other hand, seem to be only affected by higher salt contents (S-Tab.1). Only oospores from 5 PSU showed significant differences to 0.1 PSU (basal width p.adj. = 8.45e-3, shape p.adj. = 1.5e-4), 3 PSU (basal width p.adj. = 3.3e-4, shape p.adj. = 8.9e-8) and the reference group (basal width p.adj. = 7.7e-4, shape p.adj. = 7.9e-6). Below this salt boundary, no differences in shape or basal width can be detected. The two axes of the principal component analysis determine 96% of the cumulative variation of oospores (eigenvalue 1 = 1319.56, eigenvalue 2 = 324.37) and were determined by the characters of oospore length and oospore width (Fig. 1E).

**Figure 1.**
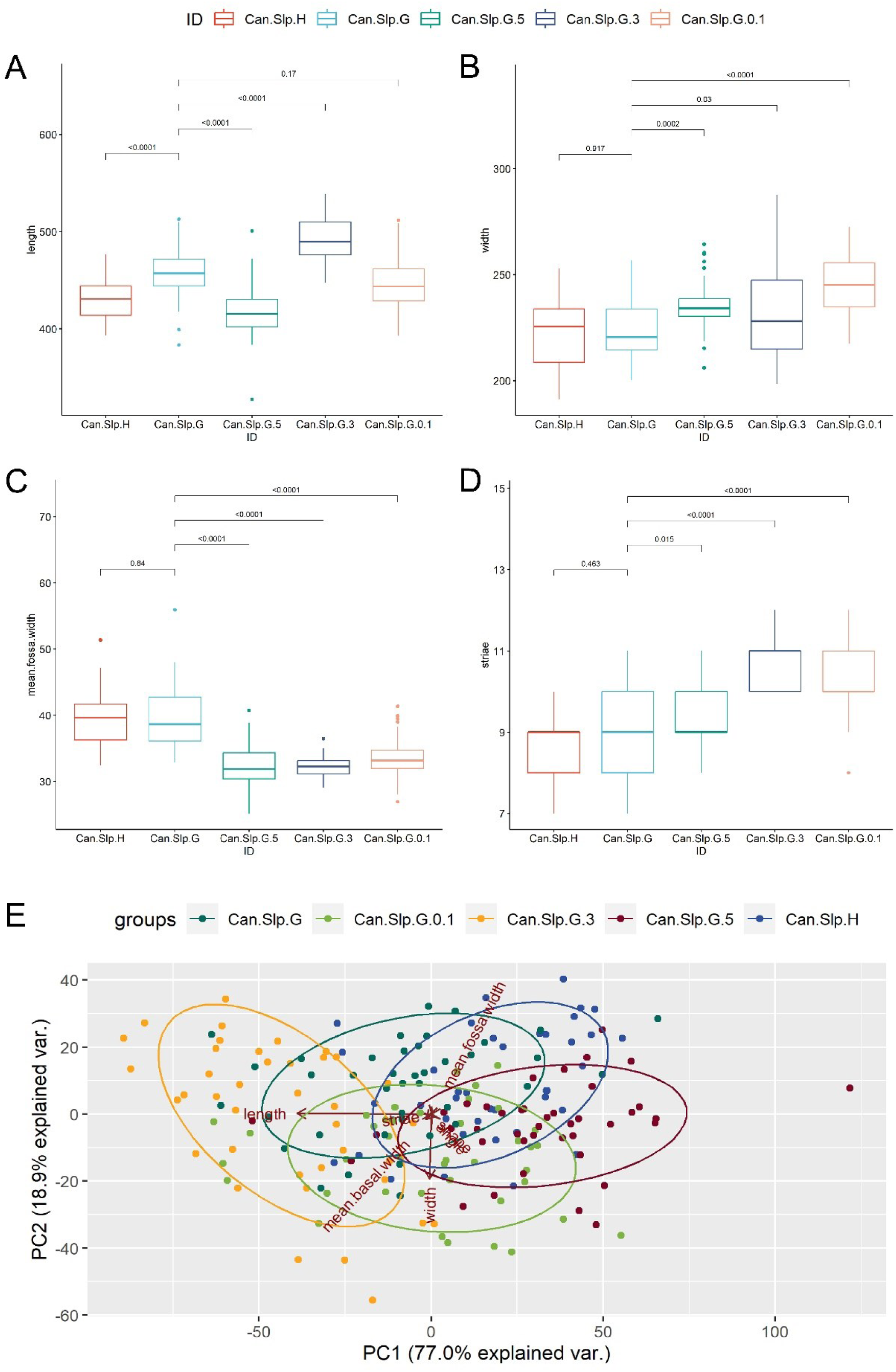
Visualisation of results for the comparison of C. canescens oospores from Neusiedler Lacken fresh material and germination experiments with different artificial salt levels. A-D. Visualisation of oospore length (A), oospore width (B), mean fossa width (C) and number of striae (D) for the five tested groups. E. Principal component analysis of oospores.

**Table 4.** Overview of morphological characters of *C. canescens* oospores from an in vitro germination experiment using different salt levels.

| variable |  | Can.Slp.G.0.1 | Can.Slp.G.3 | Can.Slp.G.5 |
| --- | --- | --- | --- | --- |
| site |  | Neusiedler Lacken<br>lower part | Neusiedler Lacken<br>lower part | Neusiedler Lacken<br>lower part |
| origin |  | in vitro germination<br>experiment | in vitro germination<br>experiment | in vitro germination<br>experiment |
| salinity (PSU) |  | 0.1 | 3 | 5 |
| n |  | 40 | 39 | 41 |
| length ($\mu\text{m}$) | min | 392.945 | 447.614 | 327.521 |
|  | max | 511.919 | 538.426 | 500.801 |
|  | mean | 448.498 | 492.224 | 416.560 |
|  | sd | 27.790 | 23.858 | 27.798 |
| width ($\mu\text{m}$) | min | 217.481 | 198.664 | 206.071 |
|  | max | 272.717 | 287.471 | 264.233 |
|  | mean | 245.177 | 232.796 | 235.227 |
|  | sd | 14.467 | 21.680 | 12.237 |
| striae | min | 8 | 10 | 8 |
|  | max | 12 | 12 | 11 |
|  | mean | 10.050 | 10.897 | 9.366 |
|  | sd | 0.986 | 0.754 | 0.915 |
| colour | RAL9005 | 100 % | 100 % | 100 % |
|  | RAL8011 | - | - | - |
|  | RAL8007 | - | - | - |
|  | sd | 0.000 | 0.000 | 0.000 |
| shape | ellipsoid prolonged | 50 % | 64.1 % | 9.8 % |
|  | ellipsoid rounded | 42.5 % | 35.9 % | 70.7 % |
|  | rounded | 7.5 % | - | 19.5 % |
|  | sd | 0.636 | 0.486 | 0.539 |
| angle($^{\circ}$) | min | 65.444 | 63.419 | 67.493 |
|  | max | 87.094 | 82.802 | 84.462 |
|  | mean | 73.994 | 73.489 | 74.796 |
|  | sd | 4.784 | 4.400 | 4.678 |
| fossa width($\mu\text{m}$) | min | 26.896 | 29.055 | 25.158 |
|  | max | 41.341 | 36.449 | 40.743 |
|  | mean | 33.537 | 32.247 | 32.530 |
|  | sd | 3.251 | 1.741 | 3.484 |
| basal width ($\mu\text{m}$) | min | 38.414 | 39.514 | 37.532 |
|  | max | 57.201 | 58.674 | 50.917 |
|  | mean | 47.775 | 49.234 | 45.340 |
|  | sd | 4.624 | 4.405 | 3.350 |
| length/width | min | 1.473 | 1.618 | 1.470 |
|  | max | 2.172 | 2.587 | 2.135 |
|  | mean | 1.836 | 2.136 | 1.774 |
|  | sd | 0.162 | 0.259 | 0.130 |

### Differences between oospores photographed with different microscopes

To exclude differences between microscopes used, oospores from Angersdorf and Neusiedler See were photographed with two different microscopes.

Anova results showed few significant differences between both microscopes and populations. Oospores from Neusiedler See Laken differ significantly in the oospore width (p.adj. = 2.3e-05), basal width (p.adj. = 6.1e-07) and striae (p.adj. = 8.3e-11), oospores from Angersdorf show differences in the oospore width (p.adj. = 5.7e-04), basal width (p.adj. =3e-03) and fossa width (p.adj. <2e-16). However, the size ranges and the results of the PCA clearly showed that the oospores of the two microscopes overlapped completely for both Angersdorf and Neusiedler See (Tab. 5). These statistical differences are caused by higher numbers of oospores photographed with microscope 1 resulting in broader size ranges (Tab. 5). Comparing mean values, no differences can be detected by use of different microscopes.

**Table 5.**
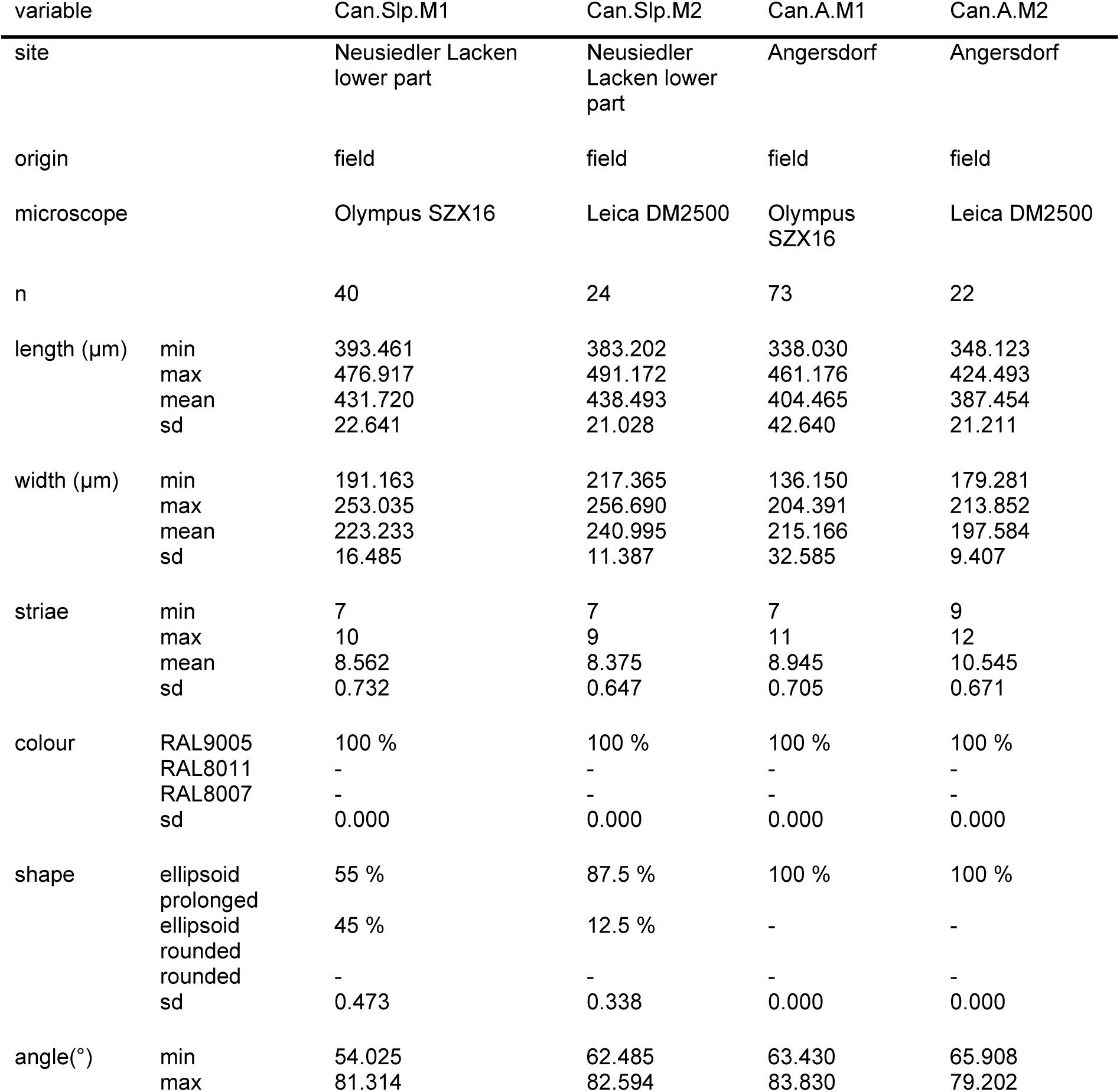
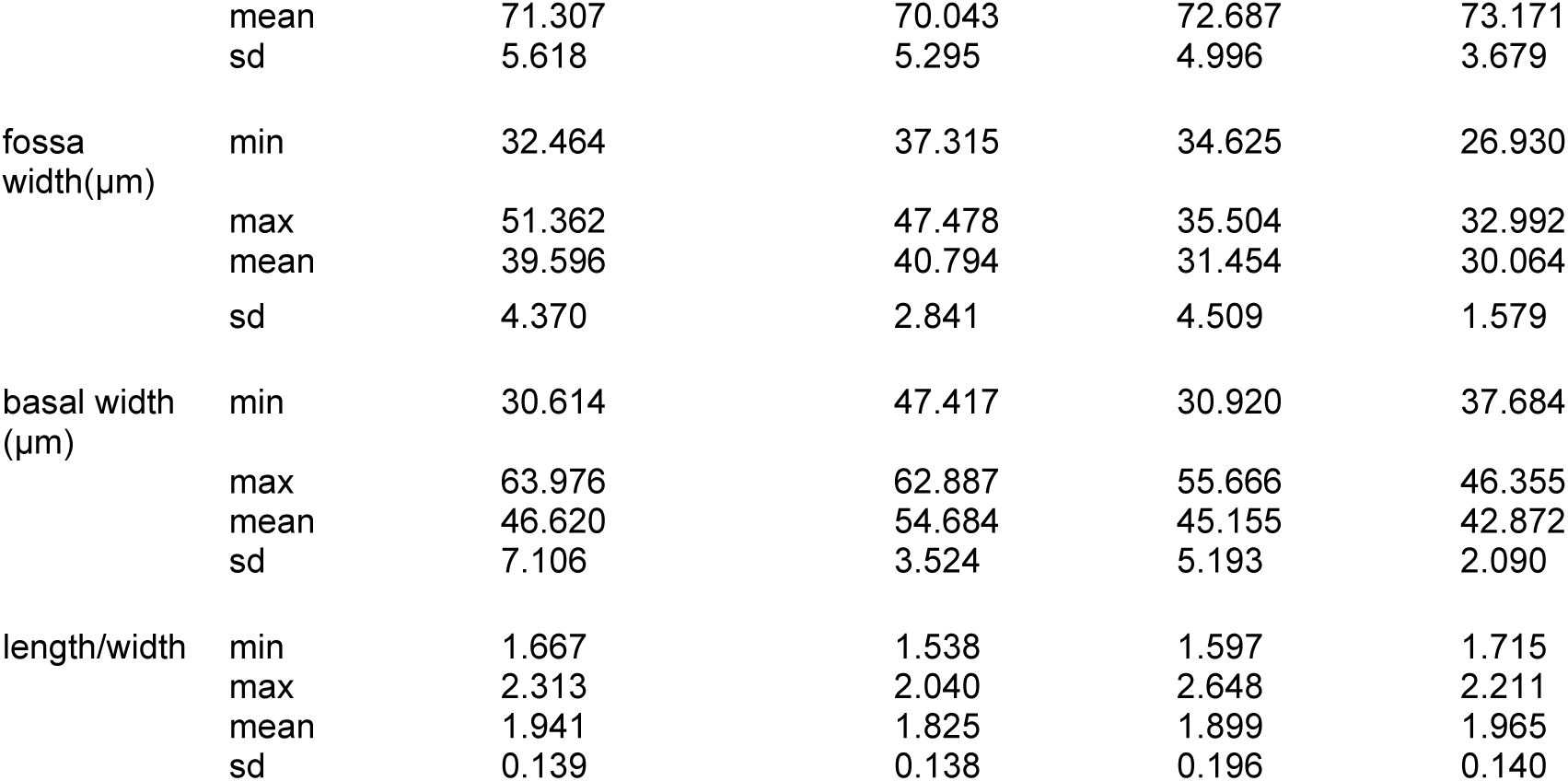
Overview of morphological characters of *C. canescens* oospores from Neusiedler Lacken and Angersdorf photographed with two different microscopes. M1 = Olympus SZX16, M2 = Leica DM2500.

### Differences between whorls

Oospores of *C. canescens* from Plóvan (France) were tested for oospore differences regarding their whorl position. For this population, no significant differences could be found that initiate position-dependent oospore sizes.

### Differences between monospecific and parthenogenetic strains

Based on the results above, two monospecific and eight parthenogenetic strains obtained from field samples were included to test whether morphological differences exist between oospores of monospecific and parthenogenetic strains. The results showed that there is no individual oospore character that allow the differentiation between both reproduction modes by oospores. However, using a model that combines morphological characters, it is possible to successfully separate any population of the data used in this study (Figure 2).

**Figure 2.**
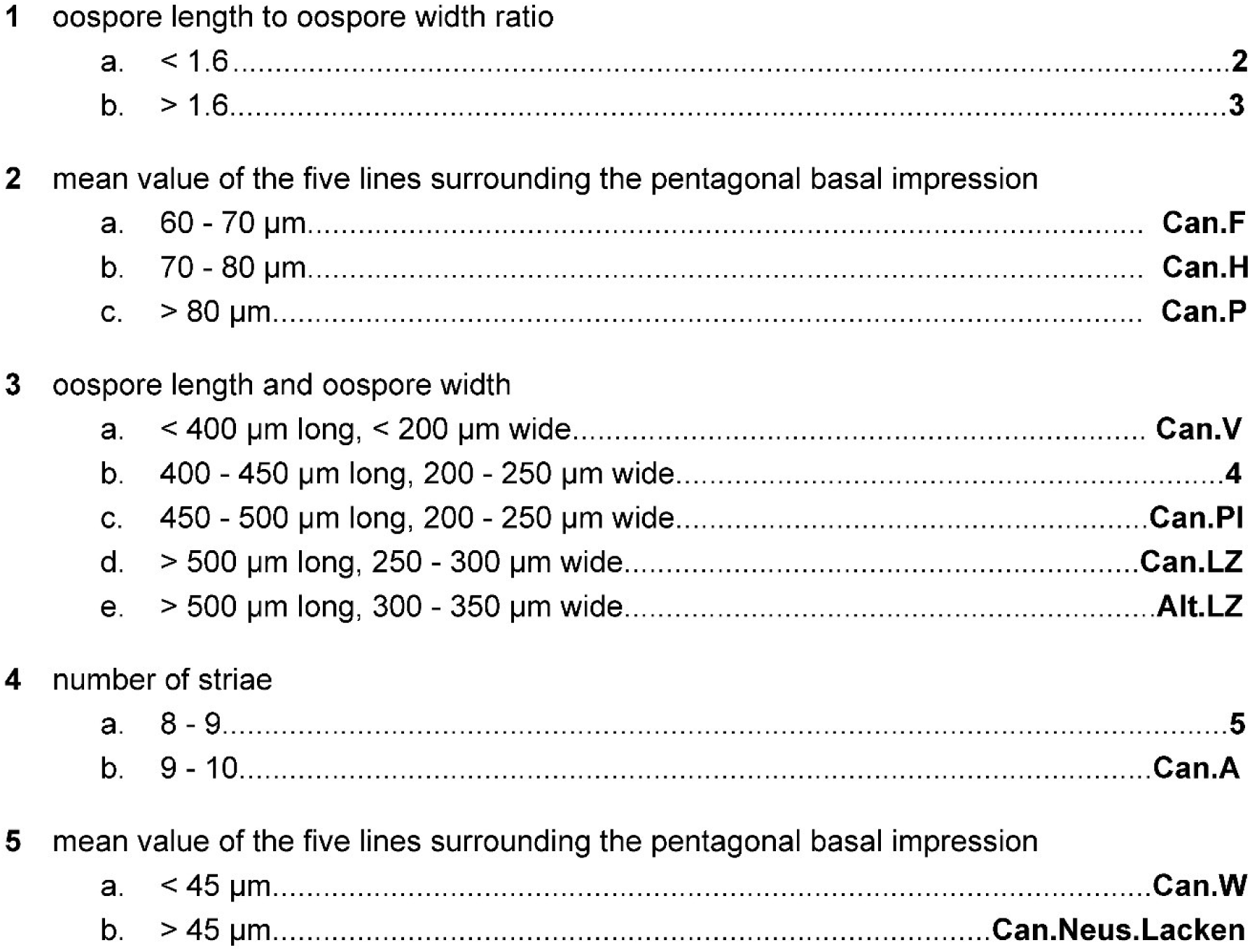
Determination key for geographically separated *C. canescens* populations of dioecious and parthenogenetic strands.

However, the two axes of the principal component analysis determine 99.2% of the cumulative variation of oospores (eigenvalue 1 = 1.18e+04, eigenvalue 2 = 1.09e+03) and were determined by the characters of oospore length and oospore width (Figure 3).

**Figure 3.**
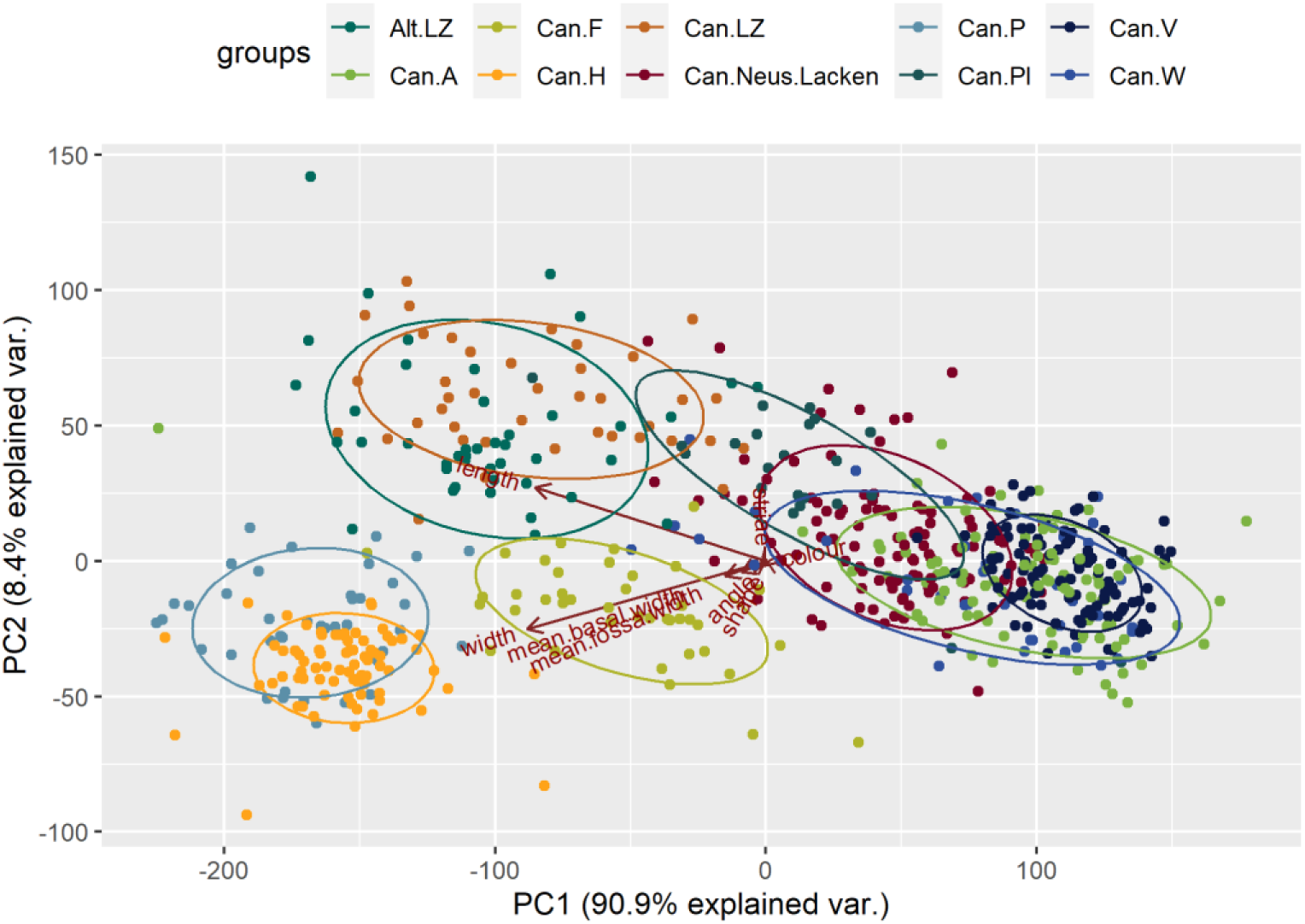
Principal component analysis of *C. canescens* populations of dioecious and parthenogenetic strands.

The ratio of these two morphological characters allow to distinguish a group of oospores from German brackish lagoons (Poel, Fastensee and Haide) from all other populations in a first step. Simultaneously, this group of oospores from German brackish lagoons could be separated by the mean value of the five lines surrounding the pentagonal basal impression. Finally, by a combination of the characters length to width ratio, oospore length and oospore width, number of striae and the mean value of the five lines surrounding the pentagonal basal impression successfully separated oospores from the Austrian dioecious population (Neusiedler Lacken) from all parthenogenetic populations.

In addition, oospores of parthenogenetic strains from marine brackish waters and lagoons (Poel, Haide, Fastensee) significantly differ from enclosed waters such as inland brackish waters (Angersdorf, Weißsee). The shape of oospores from marine conditions changes from ellipsoid prolonged to rounded (Tab.6).

**Table 6.**
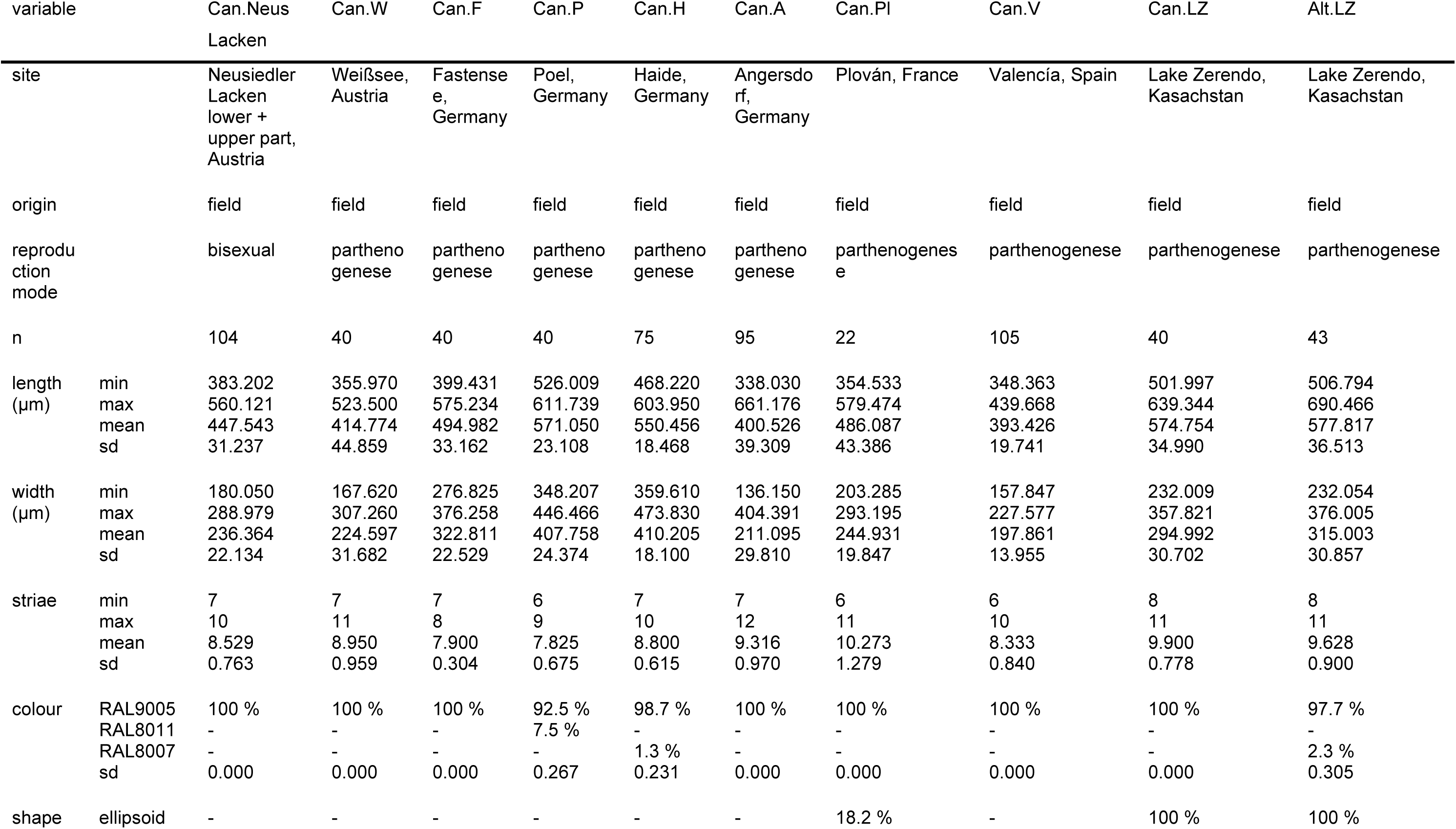
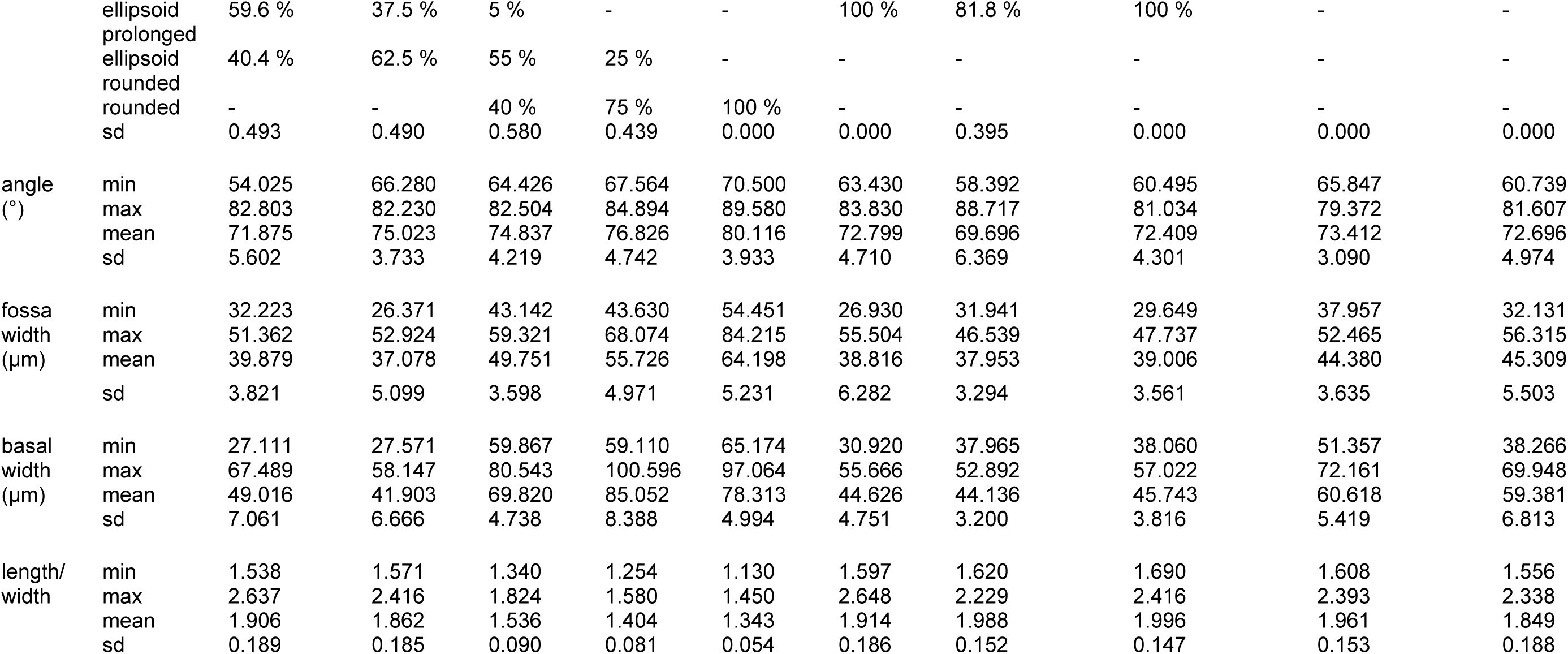
Overview of morphological oospore characters from European and Asian bisexual and parthenogenetic *C. canescens* strains as well as one Asian *Chara altaica* strain.

The oospore wall ornamentation of oospores investigated is listed in Table 7. Three European *C. canescens* populations (Austria, France and Spain) and *C. altaica* from Kazakhstan showed a pustular wall ornamentation, although differences in the shape and regularity of pustules exist. A difference between ornamentation pattern and salinity could not be found. Rather, oospores from salinity experiment showed the same sparsely pustular ornamentation whereas oospores from field samples showed a densely pustular ornamentation.

**Table 7.**
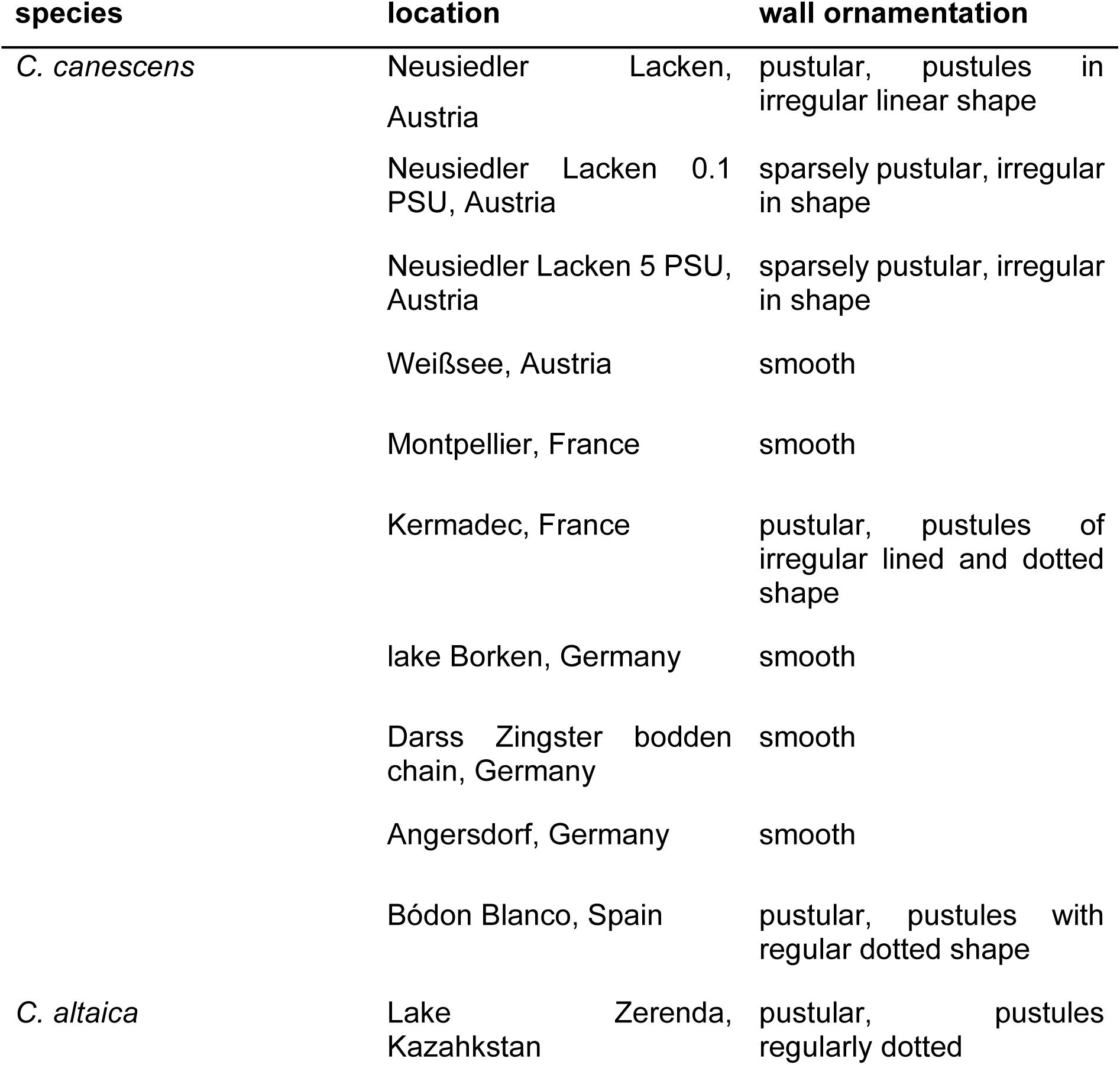
Summary of oospore wall ornamentation of *C. canescens* and *C. altaica*.

Parthenogenetic strains from Austria and Germany and herbarium sample from France showed a smooth ornamentation without pustules.

## 4. Discussion

The halotolerant species *C. canescens* is one of two European species of the section *Desvauxia* and characterised by its capability to reproduce parthenogenetically and sexually. In Europe, monoecious and dioecious populations of *C. canescens* are described (Comelles 1986, Schaible and Schubert 2008, Nowak et al. 2019), monoecious *C. altaica* populations were recently found in South Europe (Romanov and Vishnyakov *In Press*). In contrast to Europe, different monoecious and dioecious species are described for Asia and North America such as *C. hirsuta*, *C. evoluta* or *C. sibirica* (Allen 1882, 1900, Migula 1904).

Phylogenetic studies of European and Asian populations of *C. canescens* and *C. altaica* using available *rbc*L and *mat*K gene sequences from Austria, Germany, Greece, Italy, Japan, Kazakhstan, Spain and Sweden have revealed that bisexual specimens from Austria and Italy can be separated from parthenogenetic strains, *C. altaica* from Japan and Kazakhstan clustered within the group of parthenogenetically reproducing specimens from Germany, Greece, Spain and Sweden (Kato et al. 2010, Schneider et al. 2016, Langangen et al. 2019, Nowak et al. 2019).

In cases of absent vegetation or restauration purposes, taxonomic determination of oospores is performed via the stored oospore pool in sediments (Holzhausen 2018). Despite overlapping size ranges the determination of species groups is possible (Haas 1994). Furthermore, the lack of available information on oospore handling of previous studies aggravates the comparison of these data. Here we use nearly 1000 oospores sampled between 2004 and 2016, stored for re-evaluation in the Rostocker oospore-charophyte database, linked with plant material, genetics and available environmental data as well as further dry and wet stored material in a living oospore database, allowing a comprehensive regional evaluation resulting in a regional determination key for populations investigated and decisive implication for the used database.

As common for alga/plant determination keys, the use of characters combined is successfully used to differentiate the Austrian dioecious population from parthenogenetic strains. Here we used the combination of length to width ratio, the mean value of the five surrounding lines of the basal impression, the oospore length, oospore width and number of striae (Fig. 4). The oospores from Bódon Blanco are excluded. For this site, Comelles described the co-existence of the parthenogenetically and dioecious strain (Comelles 1986). Although in subsequent germination experiments only parthenogenetically reproducing alga emerged, bisexual origin could not be excluded. Comparing oospore morphology with existing phylogenetic analysis (Kato et al. 2010, Schneider et al. 2015, Langangen et al. 2019, Nowak et al. 2019), differences between oospores of parthenogenetically reproducing populations could be not reflected by sequence data of the plastid genes although the Austrian dioecious population can be separated by character combination.

**Figure 4.**
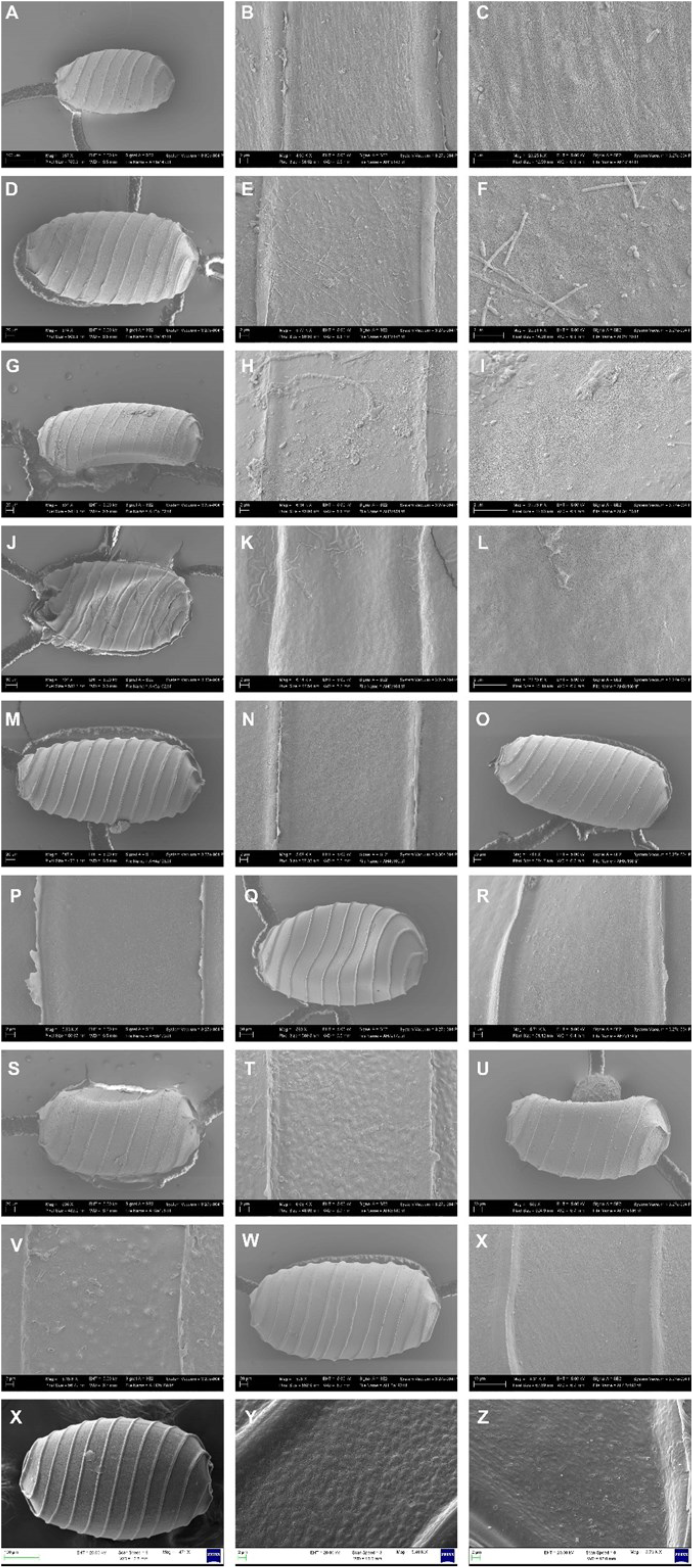
Oospores of *Chara canescens* and *Chara altaica* obtained from different populations. Listed were oospore-IDs (Holzhausen et a. 2024, *In Press*) and location. Images by M.T. Casanova. A-C Can13, Neusiedler Lacken, Austria. A) lateral view, B) fossa and ridges, C) fine details of oospore wall. D-F BB5-5N, Bodón Blanco, Spain. D) lateral view, E) fossa and ridges, F) fine details of oospore wall. G-I Can08, 0.1PSU, Neusiedler Lacken, Austria. G) lateral view, H) fossa and ridges, I) fine details of oospore wall. J-L BS56 Can02, Lake Borken, Germany. J) lateral view, K) fossa and ridges, L) fine details of oospore wall. M-N Darss Zingster bodden chain, Germany. M) lateral view, N) fine details of oospore wall. O-P Can04, Angersdorf Germany O) lateral view, P) fine details of oospore wall. Q-R Can57, Weißsee, Austria Q) lateral view, R) detail of oospore wall. S-T FR-E02 C1, Kermadec, France S) lateral view, T) details of oospore wall. U-V Can04 PSU, Neusiedler Lacken, Austria U) lateral view, V) details of oospore wall. W-X Montpellier, France W) lateral view, X) details of oospore wall. Y-AB *Chara altaica*, Lake Zerenda, Kazakhstan. Y) lateral view, Z-AB) details of oospore walls.

Furthermore, our results suppose a strong ion influence on oospore morphology features. Environmental influence on oospore morphology was hypothesized by different authors (Soulié-Märsche 1989, Casanova 1997, Holzhausen et al. 2022a, V. et al. 2022). Here, oospores of parthenogenetic populations from high fluctuating habitats as well oospores of the salt-stress-germination experiment differ from temporary populations. Combining ontogenesis and physiological studies, an accumulation of sucrose as direct storage is presumable (Holzhausen et al. 2017) increased sucrose concentrations at times of gametangia formation and maturation could explain the increased oospore volume at brackish sites with a salinity above 7.

This analysis of *C. canescens* oospores show that the oospore data base as supposed in (Holzhausen et al. 2015) and described in (Holzhausen et al. *In Press*) has the potential to elucidate regional differences in a worldwide assay.

Data on oospore origin and treatment are essential as shown by the results. Oospore sizes of field material and in vitro germination assays are comparable and could be combined consequently in analyses whereas oospores from sediment samples should be treated as separate group. Diaspore banks are known as storage pools of oospores from various previous years whereas oospores from plant material and germination assays display a time restricted period with known environmental conditions.

In contrast, our studies on oospores also have shown that no position dependency for oospores from Plovan (France) exist. The results regarding the microscope used, are in accordance with (Soulié-Märsche and Joseph 1991) who found small differences in oospores length and width for *Sphaerochara prolifera*, *S. davidii*, *Nitellopsis etrusca* and *Chara vulgaris*. In our study, these results should not be over-intellectualised, size ranges of both microscopes overlap completely. Position dependency is only known for oogonia and antheridia sizes (Ernst 1918, Calero and Rodrigo 2022). For *C. canescens* from France, no position dependency could be identified for matured oospores.

The comparison with available traits of oospores of Eurasian populations of *C. altaica*, *C. canescentiformis*, *C. evoluta*, *C. longiarticulata*, *C. pseudocanescens*, *C. piniformis*, and *C. shanxiensis* (Hollerbach and Krassavina 1983, Han and Ling 1994, Romanov 2011) yielded that all size values are mostly overlapping. Oospore surface of *C. canescens* and *C. altaica* viewed in SEM seems to be similar although variable between populations (Ray et al. 2001, Kato et al. 2010, Romanov 2011, Urbaniak 2011), plants from the same population or even within the same plant (Romanov, unpubl. data). It is described as sparsely papillate or minutely granulate to indistinctly pustular and pustular. In our study, oospores of parthenogenetic strains from Germany and Austria have shown a smooth ornamentation whereas *C. altaica* and dioecious *C. canescens* population from Austria showed a (sparsely) pustular ornamentation.

*C. canescens* oospores from France are pustular or smooth and the oospores from Spain also showed pustulars. Based on these results, only regional differences on oospore ornamentation can be fixed, a correlation between salinity and ornamentation per se do not exist. Further studies are essential to pinpoint abiotic and biotic causes for ornamentation pattering.

## Acknowledgements

The authors are thankful to A. Schoor and J. Gebert (University Rostock) for laboratory and technical assistance, C. Delker and M. Quint (MLU Halle Wittenberg) for R statistics feedback and equipment support and M. Casanova for SEM imaging. We acknowledge the constructive hints from U. Raabe (LANUV), the support and permission for sampling in protected areas: lake Borken (Egbert Korte and co-workers), French sites (Elisabeth Lambert, Didier Desmots) and Valencia (S. Romo). The laboratory equipment was partly supported by the European Fund for Regional Development (EFRD).

